# Sympatric ecological divergence with coevolution of niche preference

**DOI:** 10.1101/2019.12.20.884817

**Authors:** Pavel Payne, Jitka Polechová

## Abstract

Reinforcement, the increase of assortative mating driven by selection against unfit hybrids, is conditional on pre-existing divergence. Yet, for ecological divergence to precede the evolution of assortment, strict symmetries between fitnesses in niches must hold, and/or there must be low gene flow between the nascent species. It has thus been argued that conditions favouring sympatric speciation are rarely met in nature. Indeed, we show that under disruptive selection, violating symmetries in niche sizes and increasing strength of the trade-off in selection between the niches quickly leads to loss of genetic variation, instead of evolution of specialists. The region of the parameter space where polymorphism is maintained further narrows with increasing number of loci encoding the diverging trait and the rate of recombination between them. Yet, evolvable assortment and pre-existing assortment both substantially broaden the parameter space within which polymorphism is maintained. Notably, pre-existing niche preference speeds up further increase of assortment, thus facilitating reinforcement in the later phases of speciation. We conclude that in order for sympatric ecological divergence to occur, niche preference must co-evolve throughout the divergence process. Even if populations come into secondary contact, having diverged in isolation, niche preference substantially broadens the conditions for coexistence in sympatry and completion of the speciation process.

## Introduction

Darwin observed that many closely related species occupied the same habitat. However, he considered the sympatric origin of species by ecological divergence due to an advantage of specialists over generalists rather unlikely [1]. Since then, the contribution of sympatric speciation to observed biodiversity has remained controversial [2–5]. Speciation in a well-mixed, panmictic population is difficult for two main reasons. First, gene flow and recombination oppose divergence in polygenic traits as well as preventing reinforcement (the build-up of associations between the loci contributing to pre- and post-zygotic isolation) [6,7]. Second, the diverging types must be able to coexist. It is often thought that ecological divergence must precede the evolution of mating assortment [8], as assortment is reinforced by selection against maladapted hybrids. However, this order of events is not necessary: Assortment can precede divergence, or co-evolve with ecological loci. It is the combined effect of assortment and ecological divergence which contributes to the isolation of the nascent species [9–11]. Yet, in contrast to ecological selection, the loss of fitness due to missed heterospecific matings is typically compensated by an increase in conspecific matings. This stabilises polymorphism at the loci determining assortment and can protect polymorphism at other loci diverging under natural selection.

Habitat choice is an important source of assortment when mating occurs locally. Host-specific mating is prevalent in nature, as in phytophagous insects (reviewed in [12]). Habitat fidelity based on a preference for the hatching site is common for birds [13], another group rich in sympatric species. In general, habitat choice can be based on learned characteristics of the hatching site, on specific preference loci, or on an association with another (ecological) phenotype [14]. The first two drove the classic sympatric speciation process in the experiment by Rice and Salt [15], where fruit flies learned to choose based on phototaxis and chemotaxis. In this experiment, 60 mated females from each of two extreme habitats founded the next generation. Severe disruptive selection on multiple traits, coupled with independent regulation within equally sized niches, led to nearly complete reproductive isolation over 30 generations. Effectively, behavioural allopatry evolved (c.f. [8]). Further divergence in other traits, both selective and neutral, could have followed. While the plausibility of sympatric speciation is undisputed, how common can it really be? Selection is rarely that severe in nature, and even if niches are independently regulated, they will very rarely be perfectly symmetric.

A recognition and preference for the correct habitat is a common example of assortment by association with another phenotype. While preference for food source is omnipresent, strong [16] trade-offs in specialization of the sympatric-species-rich phytophagous insects have been hard to find, triggering a substantial controversy [17,18]. Yet, strong trade-offs must be reasonably common – otherwise, all habitats would be colonised by one generalist species. Indeed, such trade-offs have often been recovered when life-time fitness and/or performance of F1 hybrids on both hosts is assessed [19–23]. Moreover, a preference-performance correlation seems to be common in herbivorous insects [24,25].

Here we use mathematical modelling to investigate how genetically encoded preferences for the most favourable habitat facilitate sympatric speciation by ecological divergence. We assume that there is a strong (convex) trade-off in a polygenic trait, leading to disruptive selection such that generalists have a lower mean fitness than specialists. Assortment arises via preference for the correct niche. Because regulation is independent within niches (soft selection, [26]), the frequency of a type well adapted to its niche rises faster when this type is rare than when it is common. This protects the polymorphism in the ecological loci (coexistence of the specialists) via negative frequency-dependence [26].

We consider a polygenic trait under ecological divergence, and focus on assortment via preference for the niche the individual is best adapted to. The assortment allele can therefore be seen as a type of a ‘modifier-allele’ [8], realized via niche choice. The population-wide matching of the habitat thus increases with the frequency of this modifier allele. The modifier represents an ‘one-allele model’ [7], where recombination cannot break down the allele-specific association with the choice-locus. Thereby, the constraints on symmetries, which render sympatric speciation implausible, are relaxed. The modifier-allele niche choice contrasts with two-allele models, where habitat or mate choice is determined by loci independent of the ecological adaptation. Then, sympatric speciation under strong (convex) trade-off is implausible as variation is quickly lost (c.f. [7,11,27]). Such an assortment only enables evolution of coexisting specialists when trade-offs are weak (concave) [16]. Note that modifier-allele niche choice is not a fixed ‘matching’ habitat choice, where an ecological allele also directly increases the probability of getting in to the right niche (magic niche choice), which always leads to coexistence [16].

How do two nascent species evolve towards coexistence? Coexistence during divergence in sympatry is often overlooked in speciation models, side-lined by focus on growth of associations between and within polygenic traits, assuming symmetries are protected [11]. The importance of pre-existing ecological divergence and/or of pre-existing assortment for further diversification in sympatry has not been much studied. Hence, it is not clear whether divergence and coexistence in sympatry is considerably more difficult than partial divergence in allopatry or broad parapatry, followed by sympatry as assortment increases. When neutral divergence in allopatry or broad parapatry does not lead to a sufficient niche separation then (similarly to instant assortment by chromosomal variations; [28]) coexistence will not be possible. Yet, coexistence in sympatry is essential: without it, every speciation in sympatry or parapatry would lead to a subdivision of the existing range – and through time, species’ ranges would become ever smaller.

Here we aim to elucidate the importance of niche recognition in facilitating divergence under disruptive ecological selection on a polygenic trait. In other words, we study how pre-existing prezygotic isolation (niche recognition) increases and coevolves jointly with postzygotic isolation driven by a strong (convex) trade-off in adaptation to different ecological niches. Does sympatric speciation remain feasible when negative epistasis between loci increases (and hence the trade-off strengthens), and when with increasing number of loci, selection becomes weaker relative to recombination?

## Methods and results

### Model

Since we focus on the effects of disruptive selection on sympatric divergence, we use a haploid biallelic version of Levene’s [29] model with mating within niches, which is more favourable to speciation [30]. In the lifecycle of the model, individuals with non-overlapping generations are – in the basic model without assortment – randomly distributed into two niches, where selection acts. Alleles of the first type (capital letters) are beneficial in one niche and alleles of the second type (small letters) in the other niche. The individuals who survive mate within the niches and then each niche contributes to a common pool proportionally to its size. From the common pool, individuals are again randomly distributed into the two niches.

For further analysis, it is convenient to define fitness of the genotypes to be between zero and one. The presence of an allele which is deleterious in one of the niches reduces the fitness of its carrier by selection coefficient s, which is equal for all loci and both niches. The epistatic coefficient ∊ determines how much the intermediate generalist genotypes (e.g., *aB* and *Ab*) are disadvantaged relative to the specialist genotypes (e.g., *AB* and *ab* for *ϵ* < 0) and is defined as the deviation from additivity of deleterious alleles in each niche if more than one of these alleles is present. Here we are assuming a special kind of symmetric epistasis, which acts among alleles deleterious in one niche and also those in the other niche. Fitnesses of all genotypes considered in this study are defined in detail further in the text and Tables 1 – 4.

**Table 1.** Fitnesses of individual genotypes in the model without niche preference. A. Two-locus model. B. Three-locus model. Selection coefficient s determines how the deleterious alleles in niche I (a, b, c) and the deleterious alleles niche II (A, B, C) reduce the fitness relative to the well adapted genotype. The pairwise epistatic coefficient E is defined as the deviation from additivity of the two deleterious alleles present in a genotype together. We only consider ε < 0 as it generates a convex trade-off and hence disruptive selection. In the three-locus model the selection and epistasis are normalised as described above. Note that the epistatic coefficient in the intermediate genotypes in the three-locus model is present only when the genotype contains two alleles that are deleterious in the particular niche. In the other niche, the epistatic coefficient for these genotypes is absent as there is only one selected allele.

| **A** |  |  |
| --- | --- | --- |
| genotype | niche I | niche II |
| AB | 1 | $1 - 2s - \epsilon$ |
| aB, Ab | $1 - s$ | $1 - s$ |
| ab | $1 - 2s - \epsilon$ | 1 |

**Table 1. Fitnesses of individual genotypes in the model without niche preference.**
| **B** |  |  |
| --- | --- | --- |
| genotype | niche I | niche II |
| ABC | 1 | $1 - 2s - \epsilon$ |
| ABc, AbC, aBC | $1 - \frac{2}{3}s$ | $1 - \frac{4}{3}s - \frac{\epsilon}{3}$ |
| Abc, aBc, abC | $1 - \frac{4}{3}s - \frac{\epsilon}{3}$ | $1 - \frac{2}{3}s$ |
| abc | $1 - 2s - \epsilon$ | 1 |

Analytical solutions for polymorphic equilibria of our model are generally impossible to find. Instead, we have analysed the instability of the two monomorphic equilibria for allele frequencies in ecological and niche preference loci at 1 and 0. We assume that an equilibrium polymorphic at the ecological loci exists between them. This approach was used by Levene [29], Prout [31] and Hoekstra et al. [32]. Gliddon and Strobeck [33] analysed the haploid analogue of Levene’s [29] model with multiple niches, and proved that conditions for instability of the monomorphic equilibria are also necessary and sufficient conditions for stability of the unique polymorphic equilibrium. Although this conclusion is not valid for general fitnesses, such as in the presence of epistasis and/or when assortment is present, analysis of local (in)stability of the monomorphic equilibria still allows us to estimate the stability of the polymorphic equilibrium for the ecological loci in the full model for most of the parameter range. The stability is then tested numerically. We discuss a small parameter region where the local equilibrium also depends on the initial conditions. Depending on the initial allele frequencies, the system evolves either to a polymorphic equilibrium in the ecological loci, or to another equilibrium where ecological loci are fixed and the locus for assortment is polymorphic, or where all polymorphism is lost.

We complement the 2-3 locus analytical and numerical results by an individual-based haploid simulation, with up to 16 ecological loci under disruptive selection (strong trade-off), and 10 loci which gradually change the bias in the niche choice from 0 to 1. Within niches, each individual is selected at random (with replacement) to pair up, with a probability proportional to its fitness. The recombination rate r is uniform across loci, as is the mutation, set to µ = 10^−3^. Each of the N_i_ pairs has exactly one offspring, the size of niche *i* is thus kept constant at N_i_. The juveniles are then redistributed between habitats, according to their preference, which reflects linearly their number of matching alleles. For example, an A-B-c-M-M-m-[..]-m type will add to its preference 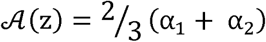, and thus choose niche 1 with a probability of 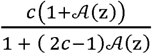, where c is the relative size of niche 1. The α sum to at most 1 over all ‘M’ loci (whilst α_i_ = 0 for all ‘m’ loci). Generations are discrete and non-overlapping.

### Ecological divergence at a polygenic trait

Ecological divergence is often driven by disruptive selection acting on polygenic traits. Therefore, we first set out to analyse how increasing the number of loci influences the ability of the model to maintain polymorphism under disruptive selection. We study two to three loci analytically and for up to 16 loci using individual-based simulations. In particular, we focus on violation of symmetries in niche proportions, which are defined as *c* = *c*_1_ =1 - *c*_11_ for niche I and II, respectively (c∊ (0,1)). In order to be able to compare the two-(*n*_*eco*_ = 2) and three-(*n*_*eco*_ = 3) locus models, we normalise the strength of selection (s) and epistasis (*∊)* in the models with more than two ecological loci such that the mean per-trait selection and epistasis remain the same (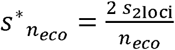and 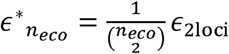). This normalisation assures that fitnesses of the specialists are always equal to 1-2s - ∊, whilst preserving the scaling of selection independently of epistasis. This leads to a slight weakening of the trade-off as the number of loci increases (see also Fig. S1).

For simplicity, we assume that all pairwise epistatic effects have the same value, and neglect higher-order interactions between alleles. Fitnesses of the individual genotypes for the two- and three-locus model are defined in Table 1A and Table 1B, respectively.

Graphical representation of the trade-offs between fitnesses of individual genotypes in niche I and niche II is shown discussed in Fig. S1 and the corresponding supplementary text.

First, we show that the region of parameter space where polymorphism is maintained is highly sensitive to violation of symmetry in niche proportions, c, even if loci are completely linked. Also, increasing convexity of the trade-off (i.e., more negative ∊, more disruptive selection) further reduces the parameter space with maintained polymorphism (Fig. 1A). Both when the loci are completely linked (r=0) and when they are freely recombining (r=0.5), there is no difference between two and three loci when mean per-trait selection and epistasis remain the same (see above). The regions of parameter space with maintained polymorphism coincide and shift towards stronger selection as epistasis increases (Fig. 1C). When recombination between loci is low (r=0.01, i.e., 1cM), the regions of parameter space with maintained polymorphism shift closer towards those of the free recombination regime in the three-locus model than in the two-locus model, as in the three-locus case the per-locus strength of selection is lower (Fig. 1B). Interestingly, there is a threshold for recombination rate, above which the parameter space where polymorphism is maintained is independent of the number of loci, which we elaborate on more in the Supplementary Information. Below we provide the analytical expressions for the boundaries of the stable regions, and the recombination threshold. The conditions are for protected polymorphism in the ecological loci, obtained by assessing local instability of the monomorphic equilibria. Note that as we have disruptive selection with symmetric selection coefficients, all ecological loci, when polymorphic, evolve to the same allele frequencies.

**Figure 1.**
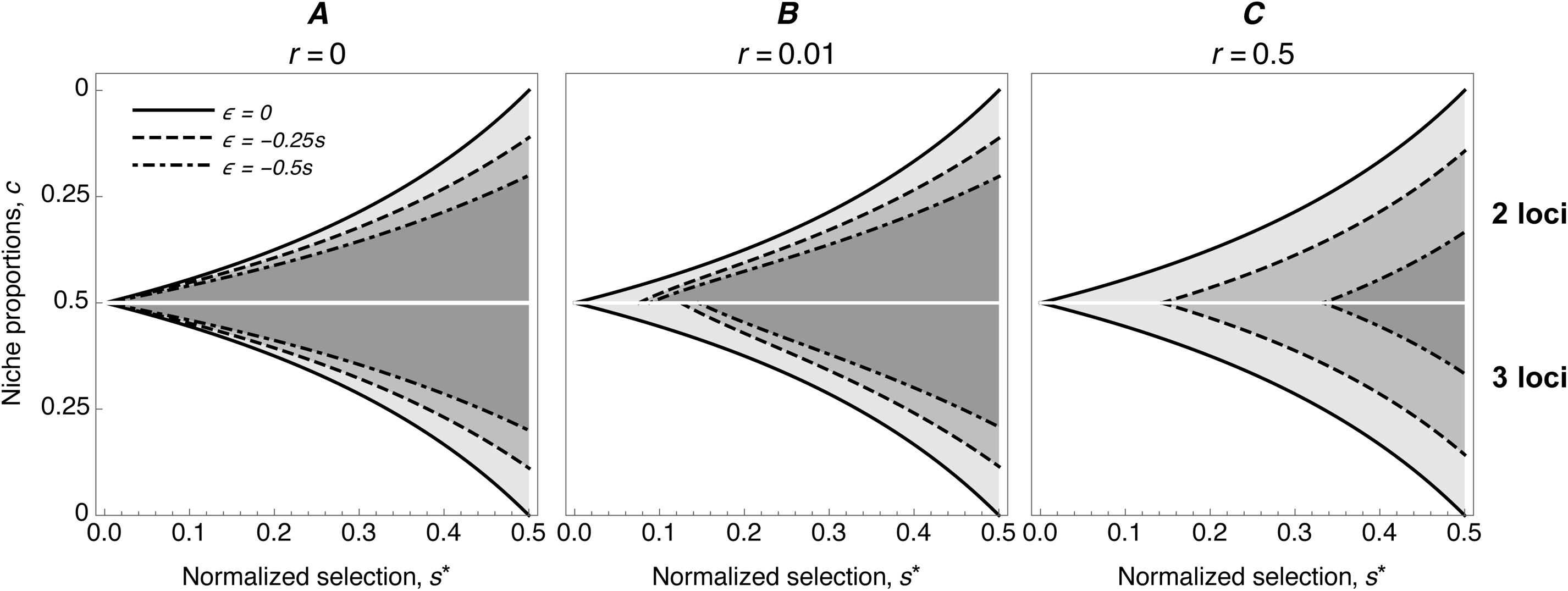
Range of niche proportions where polymorphism is maintained: where ecological divergence towards two specialists is stable. The upper half of the graphs show conditions for the two-locus model, the lower half for the three-locus model. On the x-axis is the normalized strength of selection (symmetric across loci) and on the y-axis are niche proportions. A polymorphic equilibrium is achieved for the parameter combinations of s* and *c* between the black curves and the white axis at *c* = 0.5 (shaded). Note that we are showing only one half of the parameter space for each model as the conditions are symmetrical. Therefore, every condition at a value of *c* has its symmetric counterpart at 1-*c*. The outer solid curves represent linear trade-offs (∊ = 0), the dashed curves are for (∊ = −0.25s), and the dash-dotted curves for (∊ = −0.5s). A) No recombination; B) Low recombination (r = 0.01), and C) Free recombination (r = 0.5). It is noteworthy that in the regime with low recombination (B), the region of parameter space where polymorphism is maintained decreases with increasing number of loci if selection and epistasis is normalized as described in the main text.

In both models with normalized selection and ∊ = 0 a polymorphic equilibrium is stable if

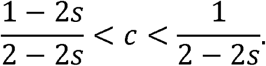

In the case with no recombination (Fig. 1A) and ∊ < 0, polymorphism is maintained if

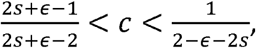

and with free recombination (Fig. 1C) if

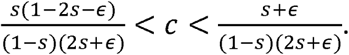

These conditions hold for the two-locus model for recombination rates 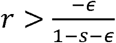. If recombination rate is low (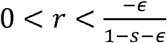; Fig. 1B), the conditions above do not hold anymore and maintenance of polymorphism is determined by

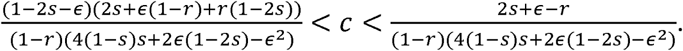

Since the equations describing the conditions for the three-locus model with low recombination and all other models presented from this point on exceed the width of a page, we confine them to the Supplementary material, where we also give more details on the stability analysis.

### Ecological divergence with niche choice

Next, we analyse how the presence of a niche preference allele influences the sensitivity of polymorphism to violation of symmetry in niche proportions. We assume that habitat choice depends both on the presence of a choosy (modifier) allele *M* and the phenotype under natural divergent selection. For the individuals with wildtype modifier allele *m*, there is no bias in selecting niches. When the modifier allele is *M*, choice depends on the selected loci in the following manner: intermediate generalist genotypes (*Ab, aB*) disperse to both niches with equal probability, and specialist genotypes (*AB, ab*) choose with probability 1 + *α* the niche which they are adapted to, and with probability 1 - *α* the niche where they are maladapted. Since in our model mating occurs within niches, this translates into a direct modification of fitnesses of the individual genotypes, as defined in Table 2.

**Table 2.** Fitnesses of individual genotypes modified by niche choice parameter α. Selection and epistasis modify fitnesses of individual genotypes as in table 1A. Genotype ABM goes to niche I with a probability increased by the factor α and to niche II decreased by α. Similarly, genotype abM preferentially goes to niche II instead of niche I.

| genotype | niche I | niche II |
| --- | --- | --- |
| ABM | $1 + \alpha$ | $(1 - 2s - \epsilon)(1 - \alpha)$ |
| ABm | 1 | $1 - 2s - \epsilon$ |
| Abm, AbM, aBm, aBM | $1 - s$ | $1 - s$ |
| abm | $1 - 2s - \epsilon$ | 1 |
| abM | $(1 - 2s - \epsilon)(1 - \alpha)$ | $1 + \alpha$ |

In the presence of niche preference, the parameter space where polymorphism is maintained broadens (Fig. 2). As allele *M* is beneficial for both specialist genotypes (*AB* and *ab*), it quickly goes to fixation when polymorphisms for the other two loci are maintained – hence, two specialists coexist. This holds in the case of complete linkage (Fig. 2A) and low recombination (Fig. 2B). In the scenario with free recombination, maintenance of polymorphism at the ecological loci depends on the initial frequencies of alleles *A, B* and *M* in the population. With weaker selection against the maladapted genotypes, s, the model is more sensitive to initial conditions, as indicated by the fading grey colour in the lower left part of Fig. 2C. Within this region, depending on the initial allele frequencies, either the preference allele M goes to fixation and protects polymorphism in the other loci, or the ecological loci go to fixation and allele M remains polymorphic and converges to its equilibrium frequency 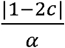. Figure 2D shows which combinations of initial frequencies of the ecological (*p*_1_ = *p*_2_) and preference (*p*_3_) alleles lead to which equilibria for four points (a, b, c, d) in the parameter space in Fig. 2C.

**Figure 2.**
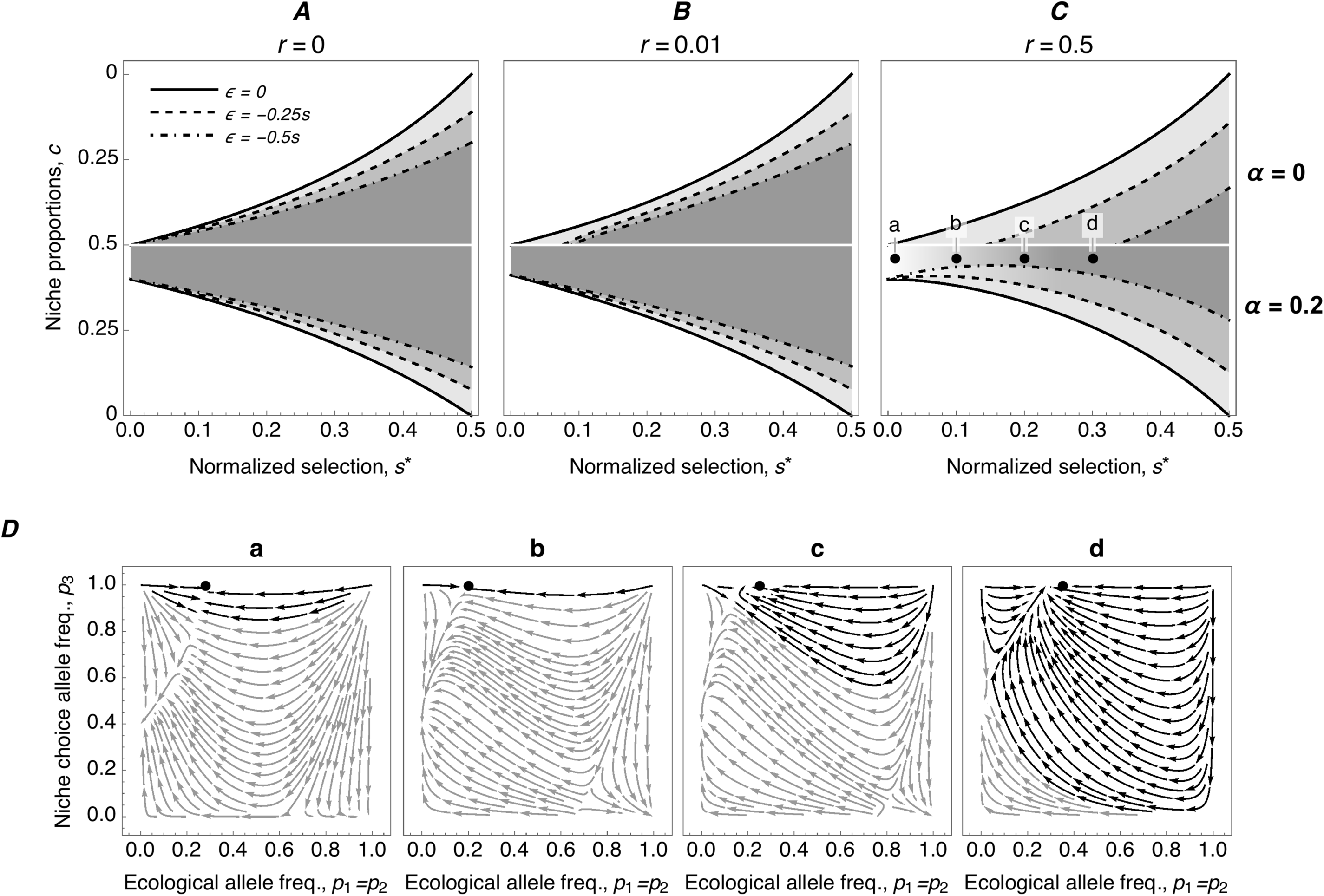
Ranges of niche proportions where ecological polymorphism is maintained (i.e., two specialists evolve and coexist), under coevolution of niche preference with ecological divergence. **A-C**: Plots showing how niche recognition allele M with niche preference *a = 0*.*2* broadens the parameter space where polymorphism at two ecological loci is maintained, relative to the model without preference (upper half of the plot; axes and all other parameters are identical to Fig 1). **D**: Streamline plots showing how maintenance of polymorphism depends on initial allele frequencies at ecological and niche preference loci for four points in Fig 2C (**a**, s=0.01; **b**, s=0.1; **c**, s=0.2; **d**, s=0.3; r=0.5, c=0.46). Black streamlines represent combinations of allele frequencies where ecological polymorphism is maintained, grey streamlines where either only ecological or all polymorphisms are lost. Since the plots were generated assuming linkage disequilibrium between ecological loci goes to 0 (i.e. loss of ecological polymorphism), the true equilibrium ecological allele frequencies, which are slightly offset with the black streamline attractors, are represented by black dots at the top of plots (**D**.**a-D**.**d**).

### Increase of assortment

Once such a niche preference modifier allele gets fixed in a population, it not only inflates the parameter space where polymorphism is maintained but it also always favours fixation of another modifier allele, which then reinforces the divergence process. In order to analyse such an increase of assortment, we again redefined fitnesses of the genotypes as shown in Table 3. The fitnesses of the specialist genotypes (*AB* and *ab*) are now defined such that the niche preference allele *M* from the previous model is fixed in the population. Therefore, both specialist genotypes have their probability of going to the right niche increased by *α*_1_. The third locus can now be polymorphic for another modifier allele, which further increases the probability of going to the right niche by *α*_2_. In Fig. 3A-C we show how the parameter space where polymorphism is maintained further broadens in the presence of another modifier allele, increasing the assortment.

**Table 3.**
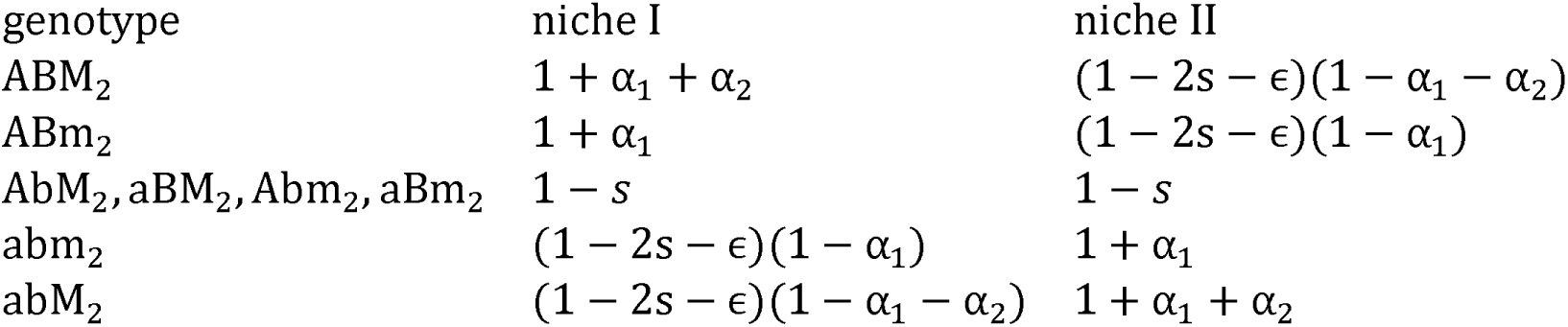
Fitnesses of individual genotypes modified by a fixed niche choice parameter *α*_1_ and a polymorphic allele M_2_, which increases niche preference for the better niche by *α*_2_. The genotype ABM_2_ goes to niche I with a probability increased by the factor *α*_1_ + *α*_2_ and to niche II decreased by *α*_1_ + *α*_2_. Similarly, genotype ABM_2_ preferentially goes to niche II instead of niche I. Preferences of other genotypes are identical as in Table 2.

| genotype | niche I | niche II |
| --- | --- | --- |
| ABM<sub>2</sub> | $1 + \alpha_1 + \alpha_2$ | $(1 - 2s - \epsilon)(1 - \alpha_1 - \alpha_2)$ |
| ABm<sub>2</sub> | $1 + \alpha_1$ | $(1 - 2s - \epsilon)(1 - \alpha_1)$ |
| AbM<sub>2</sub>, aBM<sub>2</sub>, Abm<sub>2</sub>, aBm<sub>2</sub> | $1 - s$ | $1 - s$ |
| abm<sub>2</sub> | $(1 - 2s - \epsilon)(1 - \alpha_1)$ | $1 + \alpha_1$ |
| abM<sub>2</sub> | $(1 - 2s - \epsilon)(1 - \alpha_1 - \alpha_2)$ | $1 + \alpha_1 + \alpha_2$ |

**Figure 3.**
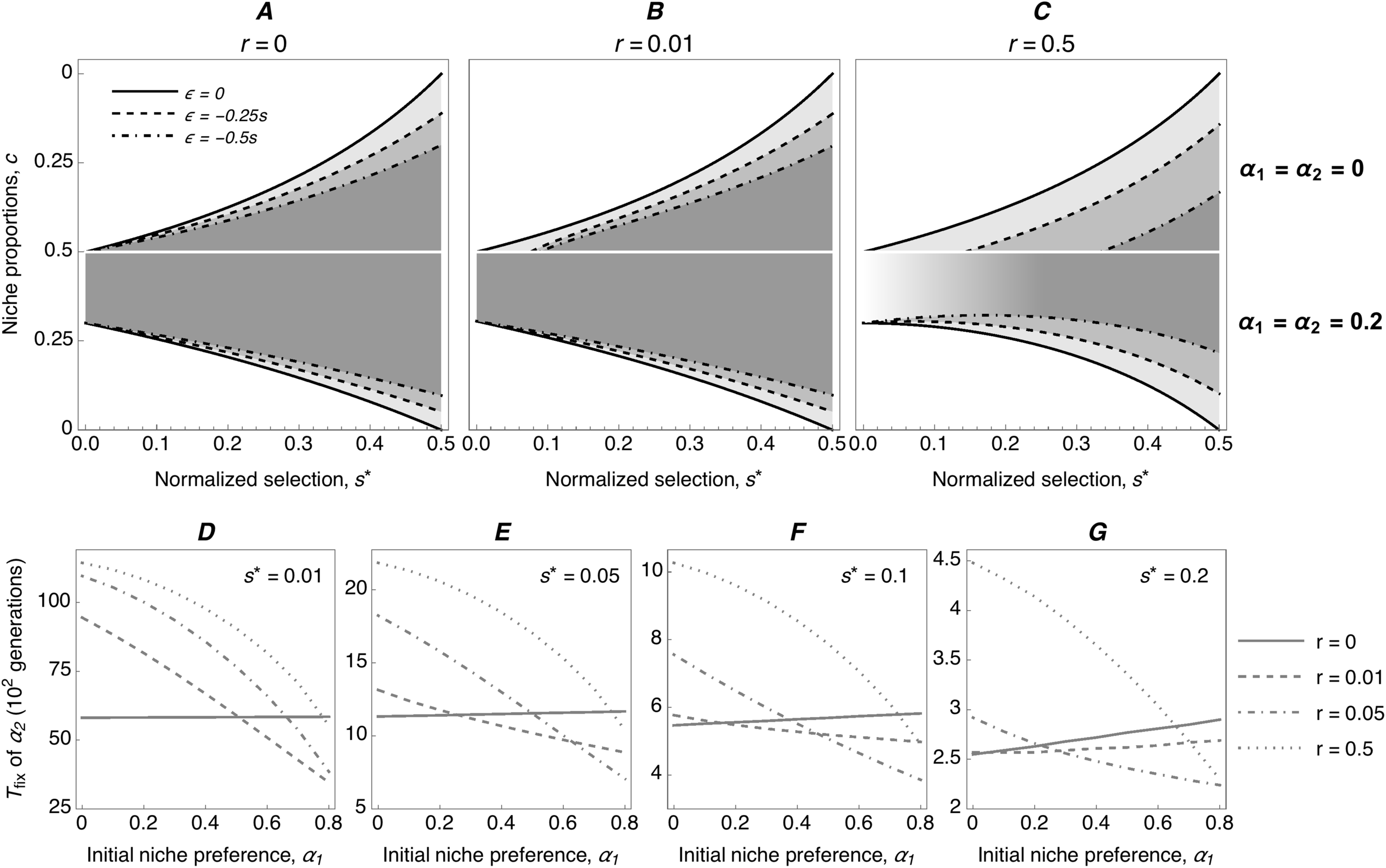
Range of niche proportions where polymorphism is maintained when assortment further increases. **A-C**: Plots showing how the parameter space where polymorphism at two ecological loci is maintained with already fixed niche preference *α*_1_ *= 0*.*2*, and an additional allele M_2_ with niche preference *α_2_ = 0*.*2*. In Figure 3C, which depicts the parameter space for free recombination (*r = 0*.*5*), the region where maintenance of ecological polymorphism depends on initial allele frequencies is indicated by a shading grey area (as in Fig. 2C). **D-G**: Time to fixation (y-axis), of the additional niche preference allele M_2_ with initial frequency *p_3_ = 0*.*01* for various combinations of initial niche preference, *α*_1_ (x-axis), strengths of selection, *s*^*^, and recombination rates, *r*. All plots are for the same disruptiveness of selection, *∊ = −0*.*5s*. The time to fixation is reduced most when selection is weak (*s*^*^ *= 0*.*01, 0*.*05*), and recombination rate is low (*r = 0*.*01, 0*.*05*) (**D, E**). In contrast, with moderate or strong selection (*s*^*^ *= 0*.*1, 0*.*2*) and low recombination (*r = 0*.*01*) the time to fixation of the niche preference allele M_2_ either decreases very little or even slightly increases with increasing initial niche preference, *α*_1_. Note that the values on the y-axis (time to fixation) differ in D-G.

Furthermore, we numerically tested how different levels of pre-existing niche preference affect time to fixation of a new niche preference allele. In Fig. 3D-G we show that with increasing initial niche preference (a_1_) the time to fixation of a new niche preference allele (*α*_2_ = 0.2) decreases. In particular, this effect is most profound when selection (s*) is weak and recombination (*r*) is low (Fig 3D, E).

### Secondary sympatry

It should be noted that all conditions presented above are independent of the initial linkage disequilibrium between the ecological loci. Therefore, for maintenance of genetic polymorphism in the ecological loci it is irrelevant whether the population starts in a Hardy-Weinberg linkage equilibrium or if it consists of only two specialist genotypes as it would during a secondary contact after divergence in allopatry.

### Cost of niche choice

It has been suggested by de Meeûs et al. [34] that in a model with soft selection and habitat preference, the conditions under which polymorphism is maintained appear to be slightly broadened but only if there is no cost to habitat selection. We implemented a cost of niche choice, *γ*, into our model, which is defined as a fraction of the selection coefficient (fitnesses are described in Table 4). The cost is associated with presence of the allele that allows for niche discrimination and choice, therefore, it is present even if the individual genotype has no preference for any of the niches but carries the niche preference allele *M*. In other words, it can be seen as a cost of an ability to discriminate (and choose).

**Table 4.** Fitnesses of individual genotypes modified by a fixed niche choice parameter *α*_1_ a polymorphic allele M_2_, which increases niche preference for the better niche by *α*_2_, and a cost of this allele, *γ*.

| genotype | niche I | niche II |
| --- | --- | --- |
| ABM<sub>2</sub> | $1 + \alpha_1 + \alpha_2 - \gamma$ | $(1 - 2s - \epsilon)(1 - \alpha_1 - \alpha_2 - \gamma)$ |
| ABm<sub>2</sub> | $1 + \alpha_1$ | $(1 - 2s - \epsilon)(1 - \alpha_1)$ |
| AbM<sub>2</sub>, aBM<sub>2</sub> | $1 - s - \gamma$ | $1 - s - \gamma$ |
| Abm<sub>2</sub>, aBm<sub>2</sub> | $1 - s$ | $1 - s$ |
| abm<sub>2</sub> | $(1 - 2s - \epsilon)(1 - \alpha_1)$ | $1 + \alpha_1$ |
| abM<sub>2</sub> | $(1 - 2s - \epsilon)(1 - \alpha_1 - \alpha_2 - \gamma)$ | $1 + \alpha_1 + \alpha_2 - \gamma$ |

As analytical solutions are rather difficult to obtain, we numerically assessed how various values of *γ* affect maintenance of polymorphism in the ecological loci. In Fig. S2 we show that for low recombination (around r = 0.01), certain cost is robustly tolerated in a large region of the parameter space. E.g., in an example with *s* = 0.1, *c* = 0.46 and ∊ = −0.5*s*, cost *y* = 0.05*s* is tolerated for α_2_ ≥ 0.1, cost *y* = 0.1*s* for α_2_ ≥ 0.2, and cost *y* = 0.2*s* for α_2_ ≥ 0.3.

### Individual-based simulations: polygenic traits with more than 3 loci

We use individual-based simulations to illustrate that initial degree of assortment is essential to allow for the evolution of coupling between ecological loci. When assortment has to evolve from low frequencies, even with strong recombination (r = 0.01), selection has to be strong for the coupling to evolve (c.f. Fig. 3 for 2 ecological loci loci, approx. s > 0.1). Weak pre-existing niche preference (one locus out of ten with α = 0.1 is near fixation) allows for evolution of two coexisting specialist if linkage is strong and the number of loci is low (Fig 4 A,B – top, n_eco_ = 3 and 6). The number of loci, among which coupling evolves, increases readily with higher initial niche-recognition (Fig. 4 vs 5). For example, with two loci out of ten with α = 0.1 near fixation, variation is already maintained for a moderate number of loci (Fig 5), though coupling between the ecological loci (in colour) stays weak when recombination is high (r=0.1) (Fig 5B, bottom rows). As initial assortment increases further, conditions for coexistence broaden considerably (see Fig S4) – two nascent specialist can then diverge and coexist even for a polygenic trait under a strong trade-off.

**Figure 4.**
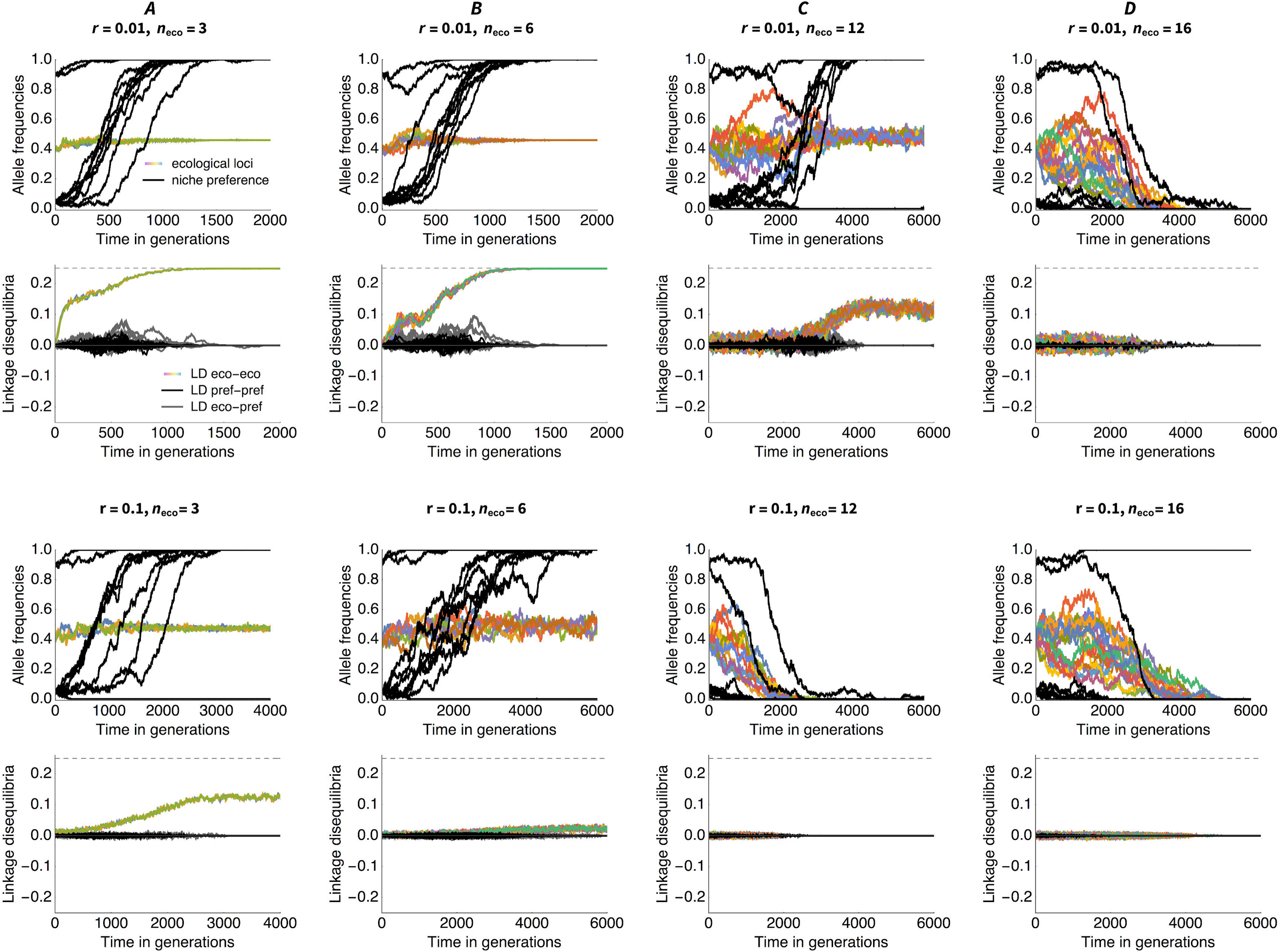
Comparison of individual based simulations with variable numbers of ecological loci (columns) and recombination between them (top two vs. bottom two rows). With moderate initial niche recognition, associations (LD) build up among multiple loci under disruptive ecological selection, and both niche recognition and ecological divergence increase jointly. In contrast, when initial niche recognition is weak, variation is quickly lost (c.f. Figure 5). First row shows the change in allele frequencies for ecological loci (in colour) and niche preference loci (black), second row shows the dynamics of the pairwise linkage disequilibria LD between ecological loci (colour), niche preference loci (black) and between ecological and niche preference loci (grey), assuming strong linkage (r = 0.01). Notice that for low number of loci (strong selection), LD starts building up immediately; whereas for higher number of loci LD only starts to build up after niche preference increases. For even higher number of loci (and/or recombination), variation is lost before assortment builds up. The quick loss of variation is facilitated by epistasis (c.f. Figure S5), genetic drift (c.f. Figure S6,S7) and recombination. Third and fourth rows depicts allele frequencies and LDs when loci are only weakly linked, r = 0.1, where joint diversification and evolution of niche preference only occurs for lower number of loci (stronger selection). Due to epistasis, loss of variation is actually faster for 12 than 16 loci. Note that both selection and epistasis are normalized with respect to the baseline two-locus case, which leads to a slight flattening of the trade-off curve as the number of loci increases (see Model and Figure S1). Parameters: s = 0.1, ε = - 0.05, slightly asymmetric niche sizes with c = 0.46, population size across niches N = 10 000. 10 niche preference loci with alpha = 0.1 (average initial niche preference for the specialist is *𝒜* = 0.22 for 2 niche-preference locus near fixation. Selection per locus is 0.1, 0.067, 0.033, 0.017, 0.0125 and pairwise epistasis is 0.05, 0.017, 0.0033, 0.00075, 0.00042 for 2,3,6,12 and 16 loci, respectively (2 loci are the reference value).

**Figure 5.**
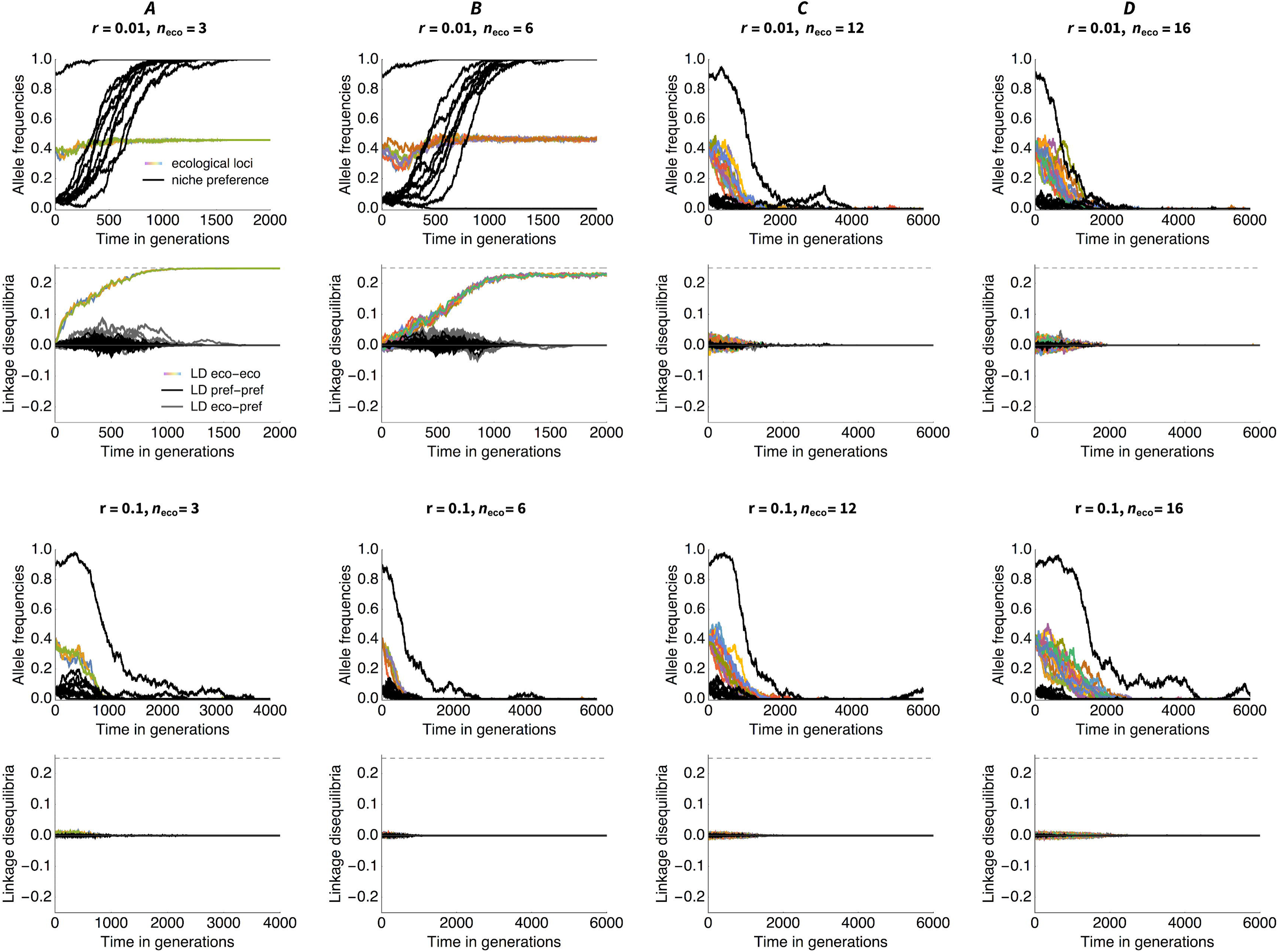
When initial niche preference is weak, divergence is only possible if ecological trait is determined by few loci, with strong selection per locus relative to recombination. With more than 6 loci (selection per locus weaker than 0.03), variation is quickly lost even when linkage is strong (r=0.01, top rows). When linkage is weak (bottom rows, r=0.1), variation is lost already for 3 loci (s = 0.67) but not 2 loci (s=0.1; not shown). Parameters are the same as in Figure 4 except here, only one of the niche-preference loci is near fixation initially, giving average initial niche preference for the specialist *𝒜* = 0.135.

For more ecological loci - assuming variation is not lost faster - we see a slow build-up of coupling between ecological loci (Fig 5C, second row). This can be seen as an abrupt phase change (tipping point, c.f. [35]). For our scenario, however, the crucial question is whether enough assortment can build up before variation is lost. Simply because the rate of change of allele frequencies grows with its heterozygosity (i.e., when their frequencies are close to ½), we see a slow increase in niche recognition initially, when its allele frequencies are low (black). It is to be noted that the niche-preference alleles are growing from low variation, not from zero – this would need different time scales (even with high mutation of μ = 10^−3^), and the waiting time does further narrow the parameter range for speciation towards coexistence. Here we focus on the later stages, assuming some variation is present.

To help the curious reader to understand the patterns in relations to other changes in the parameters, in the supplement we also include results in the absence of epistasis (when variation is maintained more easily), stronger selection (*ditto*), and a lower population size (converse). Curiously, even with total population size of N=10 000 and selection per locus s_i_ > 0.01 (16 loci), genetic drift is still important. Unfortunately, detailed exploration of these patterns is beyond the scope of this paper. We provide the source code (for Mathematica, Wolfram) and are happy to deliver further specific reference-simulations upon request.

## Discussion

Maintenance of polymorphism and the plausibility of sympatric speciation are some of the most persistent questions in evolutionary biology. An arguable assumption frequently made in many sympatric speciation models is their restriction to one or rarely several loci, whereas ecological adaptation of a population often involves a gradual change in a polygenic trait [36]. An obvious reason for this common restriction is the analytical difficulty of such models, especially if additional evolutionary forces such as epistasis and a nonlinear trade-off between viabilities are included. Notably, the multilocus version of Levene’s model – with a trade-off in adaptation to two different niches – has been analysed [37,38] but no epistasis was allowed (linear trade-off). An exception is Barton’s [39] analyses of adaptation in a quantitative trait under a strong (convex) trade-off. He assumed the symmetries in allele frequencies were maintained under disruptive and negative frequency dependent selection [40]. Yet, when niches are not perfectly symmetric, independent regulation within niches does not always generate sufficient negative frequency-dependence to stabilise ecological polymorphism. Evolution of a quantitative trait in a Levene’s model with soft selection has been also addressed using adaptive dynamics, most notably by Kisdi and Geritz [41]. They show that successive invasions and substitutions of new alleles of small effect lead to a stable polymorphism, even when niches are considerably asymmetric. Yet, branching cannot be equated with speciation. The question we are asking here is: will these branches diverge and then remain stable in the presence of recombination?

In order for specialists to be favoured over generalists in the absence of niche choice, trade-off in fitness between niches must be strong, such that a small change of the fitness in one habitat leads to a large change of fitness on the other (the fitness curve is convex). For a polygenic trait, this implies less-than-additive epistasis and/or heterozygote disadvantage. Yet, most early models of sympatric speciation focused on maintenance of polymorphism at a single diploid locus under heterozygote advantage (concave trade-off), and on coupling of such polymorphism with a locus for assortment [8,32,42]. Felsenstein [7] analysed coupling of two loci under an ecological trade-off with an independent self-recognising locus for assortment and concluded that assortment can only increase for linear or concave (weak) trade-offs. With negative epistasis (strong, convex trade-off), polymorphism was not maintained and assortment could not evolve ([7], p. 130). It has been recognised that asymmetry in niche proportions significantly influences the ability of these models to maintain genetic polymorphism – and that symmetric selection and concave trade-offs in fitness make such maintenance easier [7,8,16,26]. Curiously, Barton [39] showed that assortment can readily increase in a population under a convex trade-off, provided the starting genetic variance is large enough. However, that model assumed symmetric niches, and that symmetry in allelic frequencies is maintained by frequency-dependent selection (hypergeometric model, [40]).

We show that specialists can readily evolve even when the trade-off in fitness between niches is strong, provided that polymorphism is stabilised by assortment arising from a preference for the ‘‘correct’’ niche (to which the individual is better adapted), and there is some initial niche-recognition. As the number of loci determining the ecological trait increases, higher initial assortment is necessary to facilitate divergence. When the number of loci determining the ecological trait is small, and selection is large relative to recombination, associations (linkage disequilibria - LD) between ecological loci increase readily (c.f. [43]). With a higher number of loci (weaker per locus selection), initial niche recognition is essential to prevent loss of the variation driven by the negative epistasis (strong trade-off). This limits the scope for lingering before linkage disequilibrium starts building up (tipping points, as popularized by [35]). If selection is large relative to recombination, associations among ecological loci start building up straight away. As selection weakens relative to negative epistasis (i.e. disruptiveness of selection), variability gets quickly lost. Hence in the presence of epistasis, we only observe the so-called tipping points in the build-up of LD for moderate number of loci, not too strong recombination and moderate initial assortment (see Figure 4C).

In our analysis, we have neglected unprotected polymorphism. It has been shown that in Levene’s model with soft selection, as long as fitness is not a linear function of allele frequency, there are ‘extra’ unprotected equilibria, reachable only from surrounding allele frequencies [44]. Independently, Novak [45] showed that up to three polymorphic equilibria may coexist between 2 demes for 1 diploid bi-allelic locus (two of them unprotected). While we would have missed such unprotected equilibria by only testing for instability of monomorphism, our extensive numerical analysis and simulations do not suggest that these are biologically relevant in our model system (indeed, the population state space around such unprotected equilibria is expected to be very narrow [45]).

Prezygotic isolation due to pre-existing niche recognition both speeds up further evolution of niche preference and allows ecological polymorphism to increase. As it stabilises polymorphism in divergent populations under secondary contact, it is replacing the previous assortment due to geographic isolation. Similar reduction of gene flow can arise via philopatry (learned habitat preference) – but as the association with fitness is indirect, the effect is likely to be weaker [46,47].

We argue that niche recognition and preference is widespread across all life forms. Even plants’ roots have growth oriented towards nutrient-rich parts of the rhizosphere [48], motile unicellular organisms, including even bacteria, can move towards their preferred food source by chemotaxis [49,50] and among (in)vertebrates it is generally known that they can recognise and choose their preferred food or habitat. Therefore, this pre-existing ability to recognise the preferred niche may in fact “hijack” arising ecological divergence, protect it, and even further reinforce the newly arising reproductive isolation by strengthening the assortment. Furthermore, the degree of niche preference does not necessarily tightly correlate with the fitness advantage, which the niche provides to the organism. For example, bacteria and yeast tend to prefer one food source over all others (a phenomenon known as a diauxic shift), which in fact means a 100% preference even if the fitness advantage in growth rate is only minor [51,52]. In a phytophagous butterfly *Melitaea cinxia*, host plant preference has been observed but no measurable fitness advantage was detected on the more preferred host compared to the less preferred one [53]. In many other cases a preference-performance correlation has been detected [24,25]. While the genetic basis of preference remains mostly unclear, our model is consistent with many observations in ecology where animals tend to choose a diet, which is optimal for them [54].

## Acknowledgement

We would like to thank Nick Barton, Joachim Hermisson, Stuart Baird, Tiago Paixão and Tadeas Priklopil for their insightful comments on parts and/or earlier stages of this study. Also, we are grateful for many valuable comments and suggestions from the reviewers during the peer review process. PP was supported by a bilateral Czech-Austrian program Aktion, and both JP and PP were supported by Austrian National Agency (FWF P-32896B).

