## Supplementary Information for "Sympatric ecological divergence with coevolution of niche preference"

### **Supplementary material**

#### Conditions for maintenance of polymorphism used in Fig. 1B, Fig. 2 and Fig. 3.

In order to find conditions for stability of the polymorphic equilibrium we investigate the conditions for a protected polymorphism (i.e. the circumstances under which allele frequencies converge to a polymorphic equilibrium, starting from a perturbation to monomorphic equilibria). With two haploid loci, instability of both monomorphic equilibria p1 = p2 = 0 and p1 = p2 = 1 implies stability of the polymorphic fixed point (Karlin and Campbell, 1980). In general, however, there can be additional equilibria. Indeed, it has been shown that in Levene's model with soft selection, as long as fitness is not a linear function of allele frequency, there exist 'extra' unprotected equilibria, reachable only from surrounding allele frequencies (Priklopil 2012). As their population state space is very narrow (Novak 2011), the unprotected equilibria were never recovered by our numerical analysis and we neglect them throughout.

We could only find the solutions for polymorphic equilibria in the ecological loci assuming symmetric selection and free or no recombination – and for general recombination, we could not establish conditions for local stability. We have therefore analysed the behaviour of the system numerically to see when the instability of the monomorphic equilibria indeed gives us the conditions for stability of the polymorphism in the ecological loci (see esp. Figs. 2C,D a 3C).

To obtain the instability of the monomorphic equilibria, we computed the Jacobi matrices J0 and J1 of the system at 0 and 1 for the ecological loci and for 1 for the niche preference locus (i.e. 001 and 111). Now, equilibria 001 and 111 are stable if and only if all eigenvalues of J_0_ and J_1_, respectively, are in absolute value < 1. We inferred conditions for niche proportions when at least one eigenvalue is in absolute value > 1 for each of the cases treated here (i.e., r=0, r=0.01, and r=0.5). In the cases when more eigenvalues fulfilled this condition, we identified the leading (greatest) one, which gives the true conditions for instability of the monomorphic equilibrium (sometimes, this would be a function of the parameters). Note that from here on, *e* instead of$\epsilon$ is used in the conditions to determine the amount of epistasis. Also, the conditions for the transitions between low and free recombination differ in the three model types used in this study, and their derivation is explained in the section “Thresholds for transitions between conditions” below.

Polymorphism in the three-locus model under low recombination (Fig. 1B) is maintained if

$\frac{(-1+e+2s)(e{(-1+r)}^{3}+r^{2}(3-6s)-2s+r^{3}(-1+2s)+r(-3+6s))}{{(-1+r)}^{3}(e^{2}+4(-1+s)s+e(-2+4s))}<c<\frac{e-3r+3r^{2}-r^{3}+2s}{{(-1+r)}^{3}(e^{2}+4(-1+s)s+e(-2+4s))}$.

For the two-locus model with the third locus encoding habitat preference (Fig. 2) these conditions are the following:

For Fig. 2A:

$\frac{(-1+e+2s)(-1+\alpha)(e+2s+\alpha)}{e^{2}(-1+\alpha)-4s(-1+\alpha)+4s^{2}(-1+\alpha)+2e(-1+2s)(-1+\alpha)+2\alpha}<c<\frac{e+2s+\alpha+\alpha^{2}-e\alpha^{2}-2s\alpha^{2}}{e^{2}(-1+\alpha)-4s(-1+\alpha)+4s^{2}(-1+\alpha)+2e(-1+2s)(-1+\alpha)+2\alpha}$,

for Fig. 2B:

$\frac{(-1+e+2s)(-1+\alpha)(e(-1+r)(-1+\alpha)+r(-1+2s)(-1+\alpha)+2(s+\alpha-s\alpha))}{(-1+r)(4(-s{(-1+\alpha)}^{2}+s^{2}{(-1+\alpha)}^{2}-\alpha)+e^{2}{(-1+\alpha)}^{2}+2e(-1+2s){(-1+\alpha)}^{2})}<c<-\frac{(1+\alpha)(r-2s+e(-1+\alpha)-2\alpha+r\alpha+2s\alpha)}{(-1+r)(4(-s{(-1+\alpha)}^{2}+s^{2}{(-1+\alpha)}^{2}-\alpha)+e^{2}{(-1+\alpha)}^{2}+2e(-1+2s){(-1+\alpha)}^{2})}$,

and for Fig 2C:

$\frac{(-1+e+2s)(-1+\alpha)(s+\alpha)}{(-1+s)(e(-1+\alpha)+2s(-1+\alpha)-2\alpha)}<c<-\frac{(1+\alpha)(e(-1+\alpha)-\alpha+s(-1+2\alpha))}{(-1+s)(e(-1+\alpha)+2s(-1+\alpha)-2\alpha)}$.

For the extended niche preference model (Fig. 3) the conditions where polymorphism is maintained are:

For Fig. 3A:

$\frac{(-1+e+2s)(-1+\alpha1+\alpha2)}{2+e(-1+\alpha1+\alpha2)+2s(-1+\alpha1+\alpha2)}<c<\frac{1+\alpha1+\alpha2}{2+e(-1+\alpha1+\alpha2)+2s(-1+\alpha1+\alpha2)}$,

for Fig. 3B:

$\frac{(-1+e+2s)(-1+\alpha1+\alpha2)(e(-1+r)(-1+\alpha1+\alpha2)+r(-1+2s)(-1+\alpha1+\alpha2)+2(\alpha1+\alpha2-s(-1+\alpha1+\alpha2)))}{(-1+r)(e^{2}{(-1+\alpha1+\alpha2)}^{2}+2e(-1+2s){(-1+\alpha1+\alpha2)}^{2}+4(-\alpha1-\alpha2-s{(-1+\alpha1+\alpha2)}^{2}+s^{2}{(-1+\alpha1+\alpha2)}^{2}))}<c<-\frac{(1+\alpha1+\alpha2)(e(-1+\alpha1+\alpha2)+2s(-1+\alpha1+\alpha2)-2(\alpha1+\alpha2)+r(1+\alpha1+\alpha2))}{(-1+r)(e^{2}{(-1+\alpha1+\alpha2)}^{2}+2e(-1+2s){(-1+\alpha1+\alpha2)}^{2}+4(-\alpha1-\alpha2-s{(-1+\alpha1+\alpha2)}^{2}+s^{2}{(-1+\alpha1+\alpha2)}^{2}))})$,

and for Fig. 3C:

$1+\frac{(1+\alpha1+\alpha2)(-\alpha1-\alpha2+e(-1+\alpha1+\alpha2)+s(-1+2\alpha1+2\alpha2))}{(-1+s)(e(-1+\alpha1+\alpha2)+2s(-1+\alpha1+\alpha2)-2(\alpha1+\alpha2))}<c<-\frac{(1+\alpha1+\alpha2)(-\alpha1-\alpha2+e(-1+\alpha1+\alpha2)+s(-1+2\alpha1+2\alpha2))}{(-1+s)(e(-1+\alpha1+\alpha2)+2s(-1+\alpha1+\alpha2)-2(\alpha1+\alpha2))}$.

#### Normalisation of fitness and epistasis

As described in the main text, in order to be able to compare the model with two ecological loci with models with more loci, we normalise the strength of selection ($s$) and epistasis ($\epsilon$) such that the mean per-trait selection and epistasis remain the same (${s^{*}}_{n_{eco}}=\frac{2s_{2\mathrm{loci}}}{n_{eco}}$ and ${\epsilon^{*}}_{n_{eco}}=\frac{1}{\binom{n_{eco}}{2}}\epsilon_{2\mathrm{loci}}$). This normalisation ensures that fitnesses of the specialists are always equal to $1-2s-\epsilon$ and 1. If we plot trade-offs in fitnesses of individual genotypes for all models considered in this study (Fig. S1), we find that they do not lie on the same trade-off curve defined by the following equation:

$$w_{2}=\theta+\frac{(\theta-1)(w_{1}-1)}{1+w_{1}(\beta^{2}-1)-\beta^{2}\theta}.$$

With increasing number of loci, the trade-off is converging from a curve with $\beta=2$ towards $\beta=\sqrt{2}$. While it may seem that the strength of disruptive selection is decreasing with increasing number of loci, we are convinced that in fact the disruptiveness of selection remains the same because the effect of disruptive selection is only distributed across more genotypes. In other words, with decreasing number of loci the effect is only more and more discretised. But irrespective of the number of encoding loci, the alleles encoding the maladapted phenotype are experiencing the same overall disadvantage. As a result, in all cases the mean fitness across all possible phenotype values for any number of loci in each niche remains equal to $1-s-\frac{e}{3}$. Therefore, if the loci are recombining freely or not at all, there is no difference in the behaviour of the two and three locus model (with r=0 they behave like a single locus and with r=0.5 all alleles are in linkage equilibrium in the absence of assortment). However, when recombination is low enough but appreciable (e.g., r=0.01), the associations between beneficial alleles break up faster with higher number of alleles, as the overall average recombination rate per trait becomes higher. As a result, the recombination rate threshold for maintenance of polymorphism decreases with increasing number of loci encoding the selected trait, i.e., with higher number of loci, lower recombination rate is “tolerated” by disruptive selection in order to maintain polymorphism as LD breaks down faster than in a model with lower number of loci.

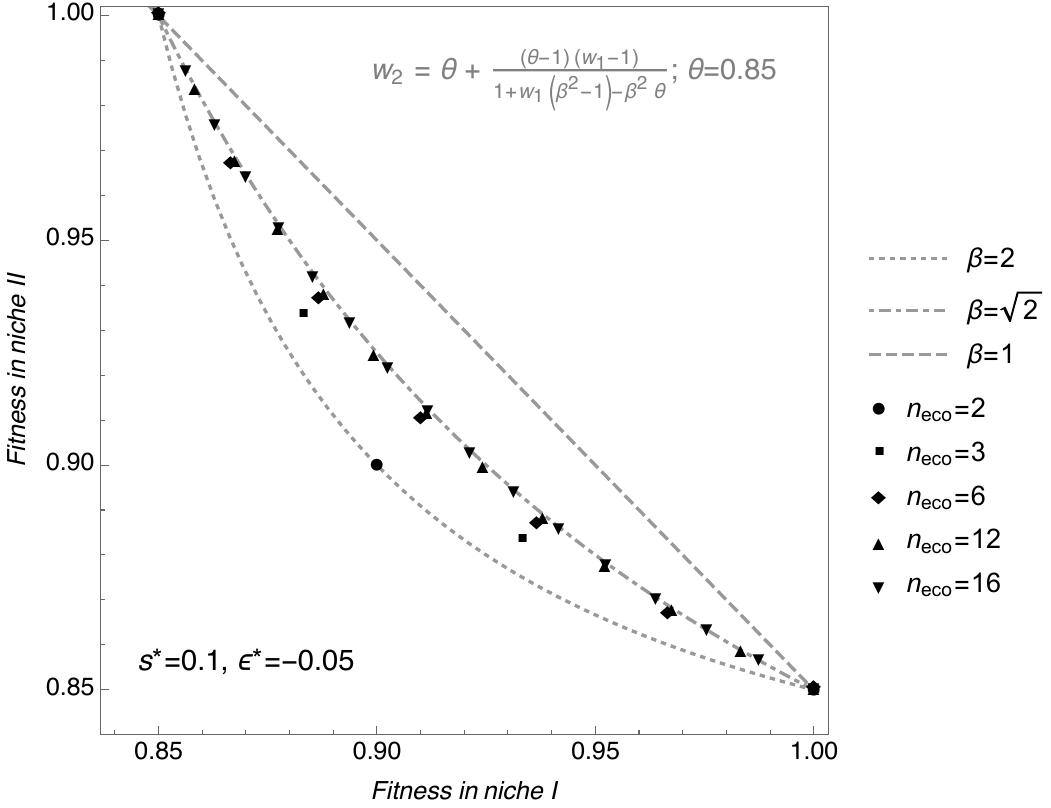

**Figure S1.** Trade-offs between fitnesses in niche I and niche II.

Grey curves show functional representation of fitnesses with different parameters of $\beta$. Grey dashed line ($\beta=1$) represents linear trade-off, grey dot-dashed curve represents convex trade-off with $\beta=\sqrt{2}$, and dotted grey curve with $\beta=2$. Black symbols represent individual genotypes for models with numbers of loci ($n_{eco}$) indicated in the legend.

#### Thresholds for transitions between conditions

The parameter combinations when conditions determining instability of monomorphic equilibria for the no and low recombination regime change to the conditions for the free recombination regime were determined by solving an equation when these conditions are equal for *r*. For the two-locus model, this transition happens when

$r=\frac{e}{-1+e+s}$.

In the model with three ecological loci, this transition occurs when

$r=\frac{2-4e+2e^{2}-6s+6es+4s^{2}+{((-1+s){(-1+e+s)}^{2}{(-1+e+2s)}^{3})}^{1/3}-ⅈ\sqrt{3}{((-1+s){(-1+e+s)}^{2}{(-1+e+2s)}^{3})}^{1/3}}{2-4e+2e^{2}-6s+6es+4s^{2}}$,

and in the case of the model with two ecological loci and a niche choice locus, this threshold is reached when

$r=\frac{-e+e\alpha+2s\alpha}{1-e-s+e\alpha+2s\alpha}$.

#### Cost of niche choice

We numerically evaluated a model version where the niche preference allele$M_{2}$ has a cost $\gamma$ that is defined as a fraction of the selection coefficient acting on the ecological alleles. In Figure S2, we show under what conditions the niche preference allele $M_{2}$ with varying levels of choosiness $\alpha_{2}$ and cost $\gamma$ goes to fixation depending on the level of initial (i.e., fixed, pre-existing) level of niche preference $\alpha_{1}$. While the initial niche preference $\alpha_{1}$ allows for fixation of the additional niche preference allele $M_{2}$ under a broader range of recombination rates $r$ (compare rows in each segment), its level does not have a strong effect on the fact whether or not an additional niche preference allele $M_{2}$ would go to fixation (compare columns in each segment). What determines whether the additional niche preference allele $M_{2}$ goes to fixation is mainly the level of niche preference$\alpha_{2}$ encoded by the allele $M_{2}$ and its cost $\gamma$ (compare individual segments – either mostly green or mostly blue).

Interestingly, in the example presented in Figure S2 the niche preference allele $M_{2}$ goes to fixation for a particular recombination rate $r=0.05$ even if it is lost for lower or higher recombination rates (green middle rows for $\alpha_{2}=0.05 \& \gamma=0.05s, \alpha_{2}=0.1 \& \gamma=0.1s,\mathrm{and} \alpha_{2}=0.2 \& \gamma=0.2s$. We attribute this to the fact that for the given combinations of $\alpha_{2}$ and $\gamma$ there is an optimal recombination rate which allows for sufficient recombination of the newly arisen niche preference allele $M_{2}$ into the specialist genotypes (AB and ab) while at the same time, this recombination rate does not break up association between the two ecological alleles too fast.

Furthermore, we do not consider any cost of the initial (i.e., fixed, pre-existing) niche preference allele when it is fixed, therefore, it affects all individuals in the population the same way. The cost penalises the ability of recognizing the right niche, not the process associated with the choice (e.g., the ability to taste the right food or recognize the right habitat). Moreover, we also consider the situation where there is no pre-existing niche preference, which is shown in the first column of each segment.

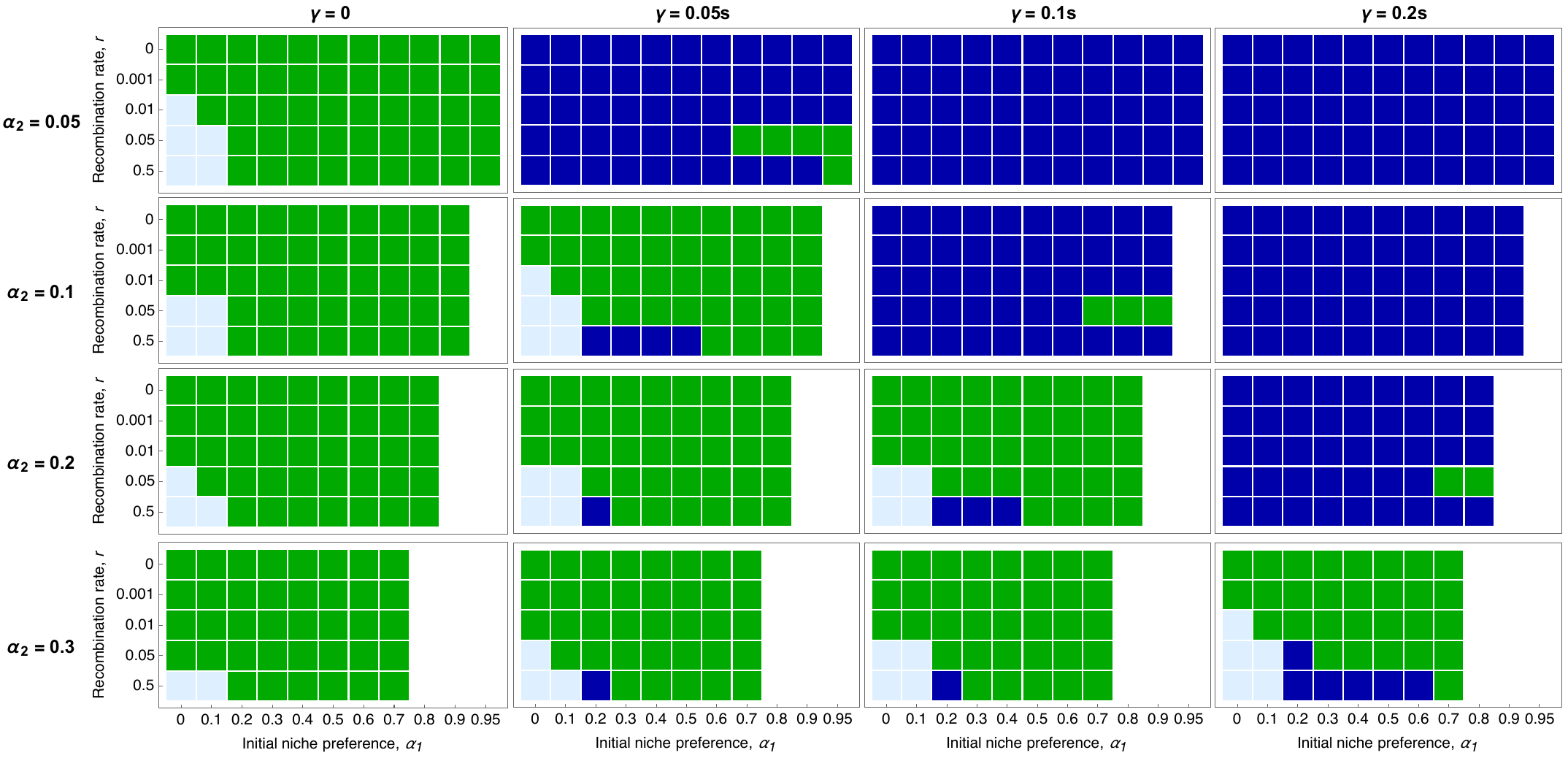

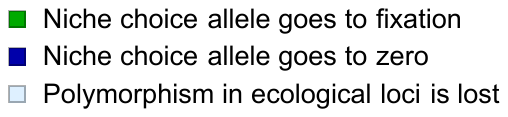

**Figure S2**. Matrix plots of parameter combinations where niche preference allele $M_{2}$ goes to fixation and polymorphism at ecological loci is maintained (green), where niche choice allele goes to zero and polymorphism at ecological loci is maintained (dark blue), and where polymorphism in both niche choice locus and ecological loci is lost (light blue). Other parameters are *s = 0.1,* *c = 0.46*, initial ecological allele frequencies *p_1_ = p_2_ = 0.5*, niche preference allele (*M_2_*) initial frequency *p_3_ = 0.01*.

#### **Dependence of polymorphic equilibria on initial niche preference allele**

We numerically assess how maintenance of ecological polymorphism is affected by initial niche preference allele frequency in the free recombination regime (Fig. 2C). For the purpose of this illustration we keep initial allele frequencies at the ecological loci constant, *p_1_=p_2_=0.5*, and are varying selection coefficient, *s*, and the initial frequency at the niche preference locus, *p_3_*. All other parameters are as in Fig. 2C *(α = 0.2, r = 0.5, ε =* – *0.5s*).

The initial niche choice allele frequency (*p_3_*) is varying from 0.01 (left, dark red) through 0.5 (grey) to 0.99 (blue). For colours of other initial *p_3_* frequencies see Fig. S1 legend. The dashed lines show the frequency of the niche preference allele (*p_3_*) after 1000 generations for varying niche proportions, *c*. The dotted line in the corresponding colour shows the frequencies of the ecological loci (*p_1_=p_2_*) after the same number of generations.

In Figure S1A we show when is polymorphism maintained, assuming selection coefficient *s = s_I_ = s_II_ = 0.05*, in Fig. S1B for *s = s_I_ = s_II_ = 0.1* and in Fig. S1C for *s = s_I_ = s_II_ = 0.2*. In all selection regimes, when *p_3_* goes to fixation (i.e. dashed line at 1), the ecological loci (i.e. dotted line) remain polymorphic. When the niche proportions, *c*, are further from symmetry, the behaviour of the model changes and *p_1_ = p_2_* go to fixation while the niche preference locus remains polymorphic. The niche preference locus remains polymorphic because when there is a high asymmetry in niche proportions (in relation to selection), it is better not to choose (the other nearly-as-good niche then stays empty). With symmetric niches, it always gives a fitness advantage to pick the right niche. It follows that by continuity, there will be a range of niche asymmetries where at least one niche-preference locus remains polymorphic

The parameter space region where *p_1_ = p_2_* go to fixation and niche preference locus remains polymorphic is larger with weaker selection (Fig. S1A, also indicated by fading grey colour in the lower left part of Fig. 2C) and is shifting to the right towards more asymmetric niche proportions with increasing strength of selection (*s = 0.1* in Fig. S1B and *s = 0.2* in Fig. S1C). Under stronger selection against maladapted genotypes in their “wrong” niche (*s*), the model more robust is the model against perturbations in niche preference allele frequencies.

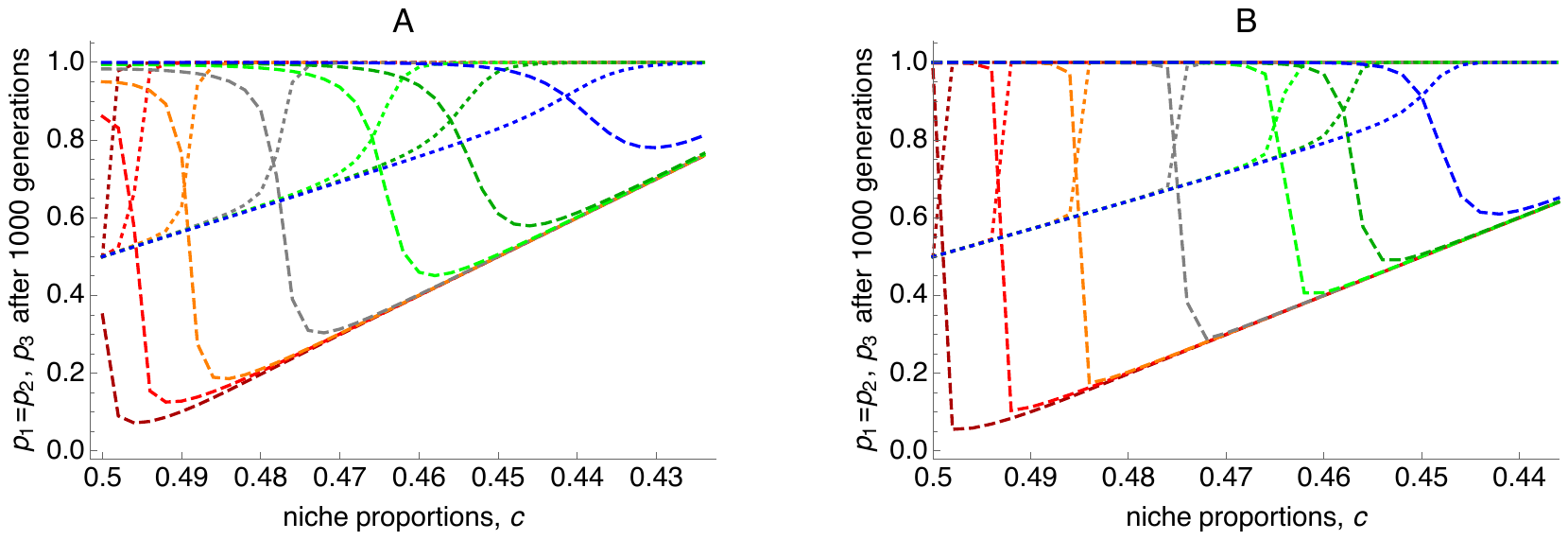

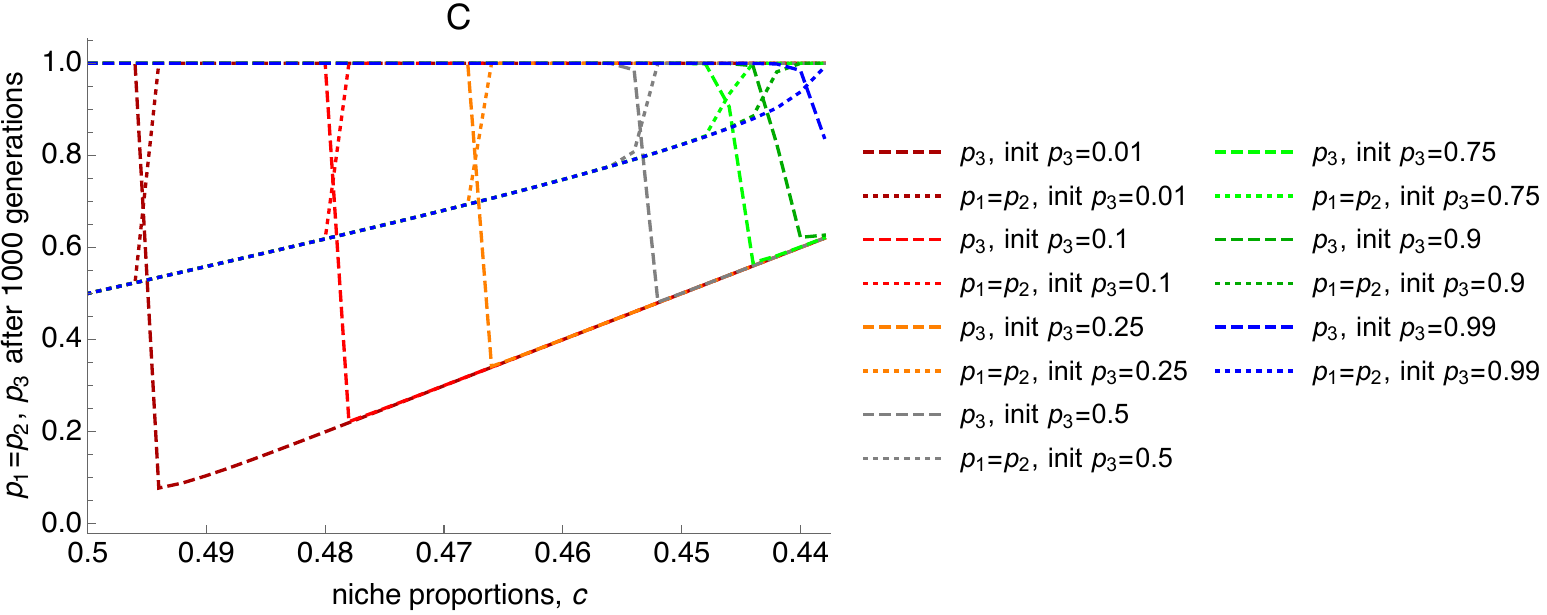

***Figure S3. Dependence of polymorphic equilibria on initial niche preference allele frequency in the free recombination regime.*** *For detailed description of the figure see the supplementary text.*

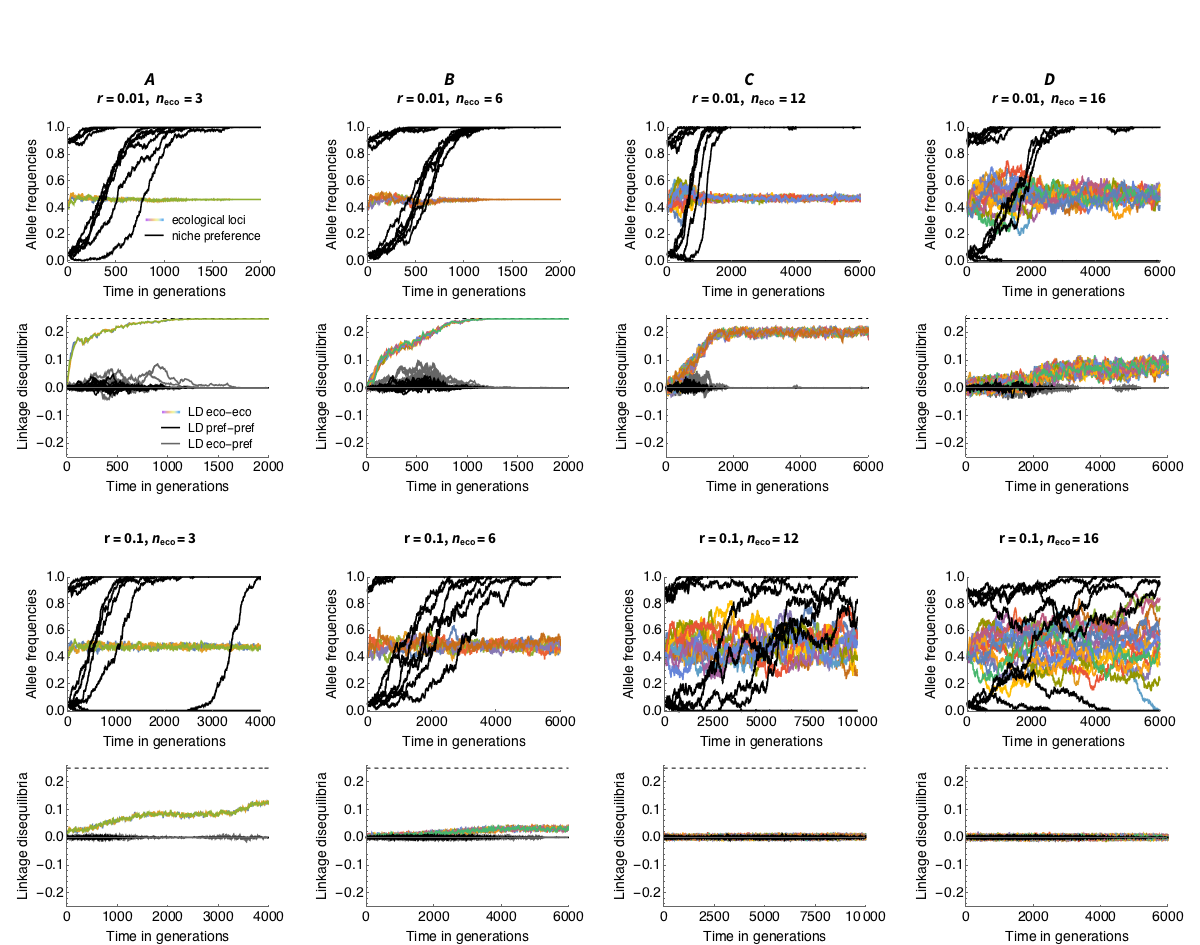

***Figure S4. With strong initial niche recognition, variation is maintained easily even under strong trade-off and weak linkage.*** *Parameters (apart from* $\mathcal{A}$ *as in Fig. 6, main text): s = 0.1, ε = - 0.05, c = 0.46, N=10.000 between niches, 10 niche preference loci with alpha = 0.1,* ***four*** *of which start at frequency of 0.9, six at frequency 0.05. This gives average initial niche preference for the specialist of* $\mathcal{A}$ *= 0.39.*

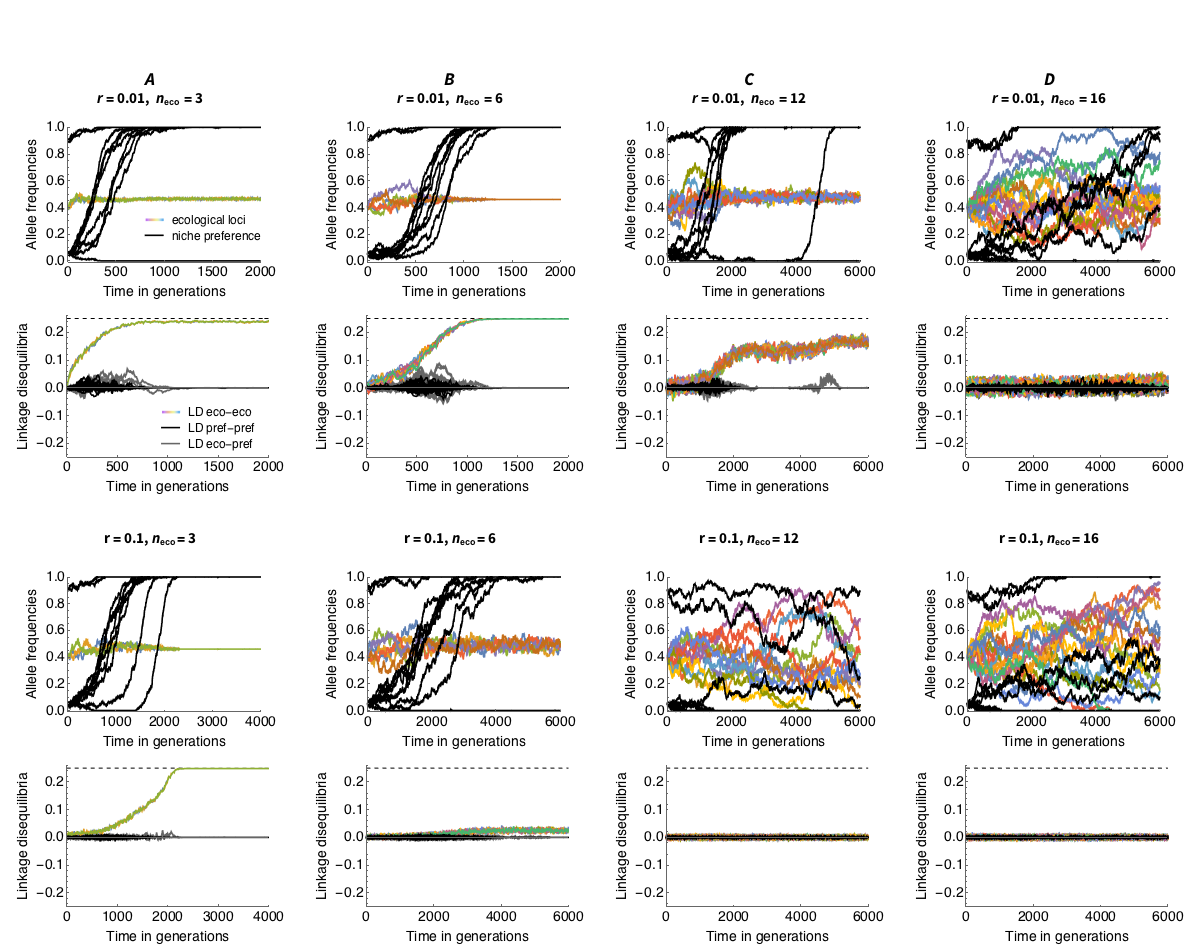

***Figure S5. In the absence of epistasis (linear trade-off), evolution towards coexistence is plausible under a broad range of parameters.*** *This is because average fitness of the generalist is the same as the average fitness of the specialist in the two niches (if they are symmetric): hence, niche preference has ample time to evolve. Parameters (as in Fig. 6, main text): s = 0.1,* ***ε = 0****, c = 0.46,* $\mathcal{A}$ *= 0.135, N=10.000.*

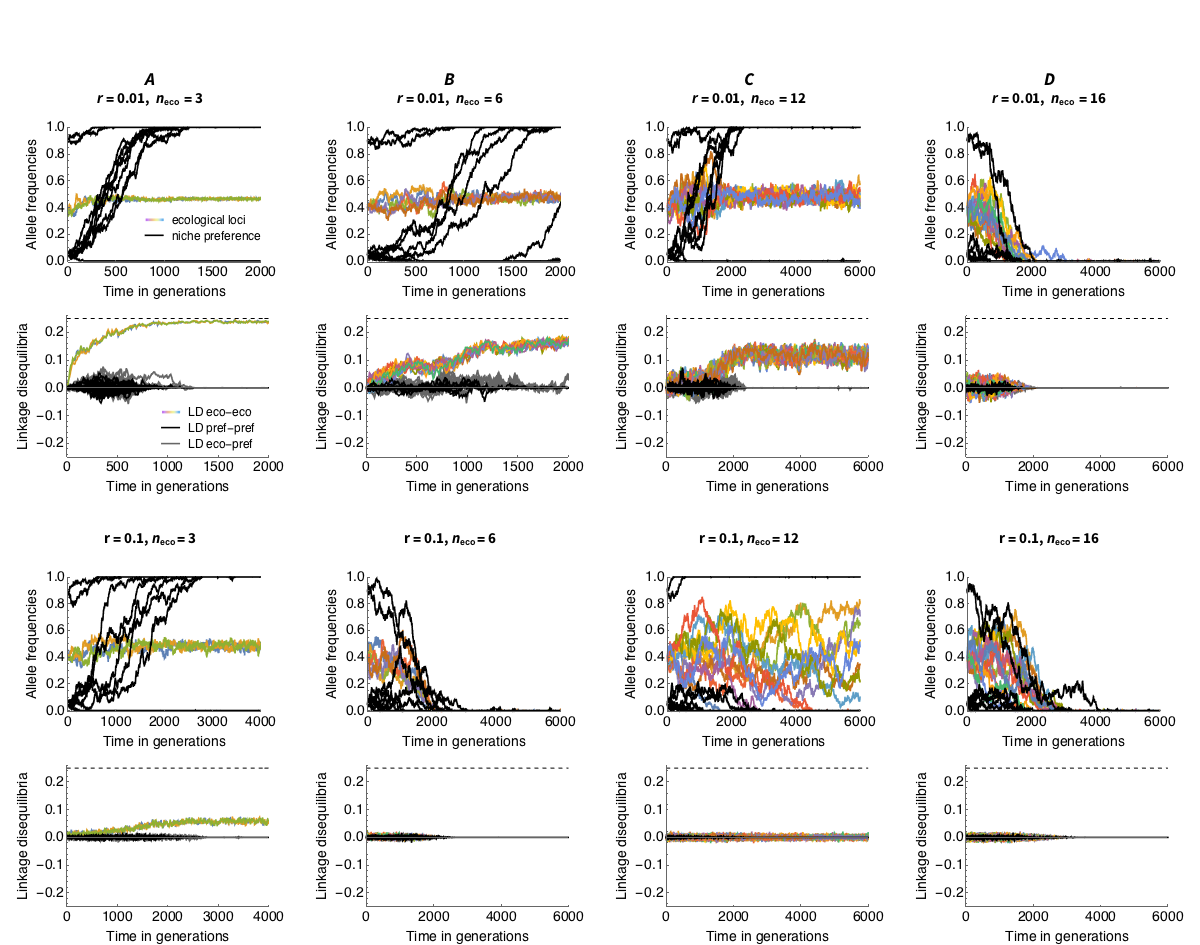

***Figure S6. Genetic drift is important and drives the fluctuations - even when niches sizes are fairly large. With a smaller population size, fluctuations are larger and variation is lost more easily.*** *Parameters (as in Fig. 6, main text): Parameters: s = 0.1, ε = - 0.05, c = 0.46,* $\mathcal{A}$ *= 0.135,* ***N=5.000****. Note that selection per locus is 0.1, 0.067, 0.033, 0.017, 0.0125 for 2,3,6 and 12 loci, respectively (2 loci are the reference value).*

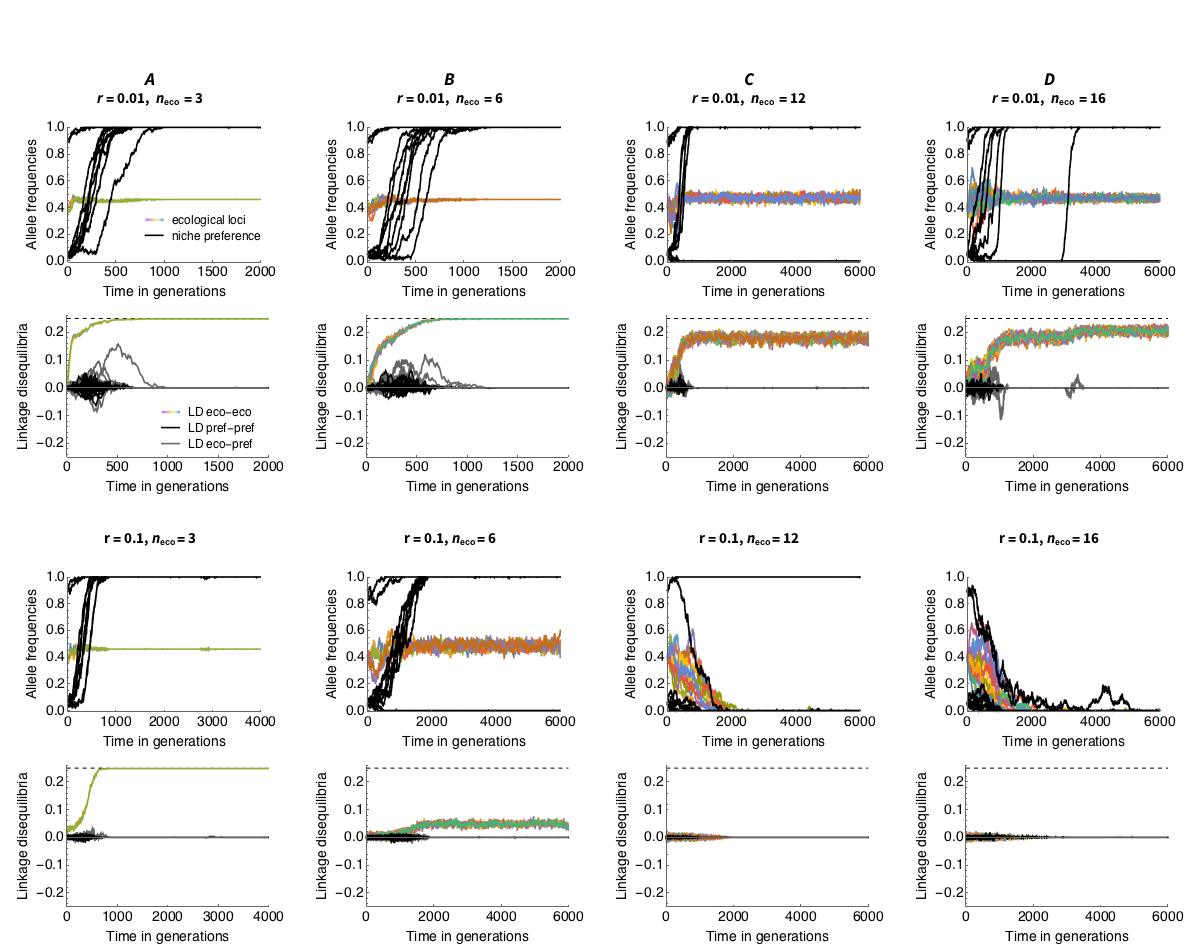

***Figure S7.*** ***Doubling selection counters the effect of genetic drift.*** *Parameters:* ***s = 0.2****, ε = - 0.05, c = 0.46,* $\mathcal{A}$ *= 0.135,* ***N=5.000****.*
